# Positive joint work redistribution in running: the role of plantar flexor fatigue and the effect of advanced footwear technology

**DOI:** 10.1101/2025.10.26.684644

**Authors:** Key Nahan, Steffen Willwacher, John Kerr, Martin Héroux, Kirsty A. McDonald

## Abstract

Over the course of a near-maximal effort 10 km run, positive mechanical work decreases at the ankle and increases at the knee. Although plantar flexor fatigue is believed to be responsible for the proximal shift in mechanical work generation, this has yet to be confirmed experimentally.

**Purpose:** To 1) determine the effect of plantar flexor fatigue on lower limb positive joint work during running and 2) determine the effect of running shoes on this relationship.

**Methods:** Trained male runners (*n* = 12) were analyzed over two 30 s runs at their 10 km race pace, without and with local plantar flexor fatigue induced via a calf-raise protocol to a level commensurate with that accumulated over an exhaustive run (∼24% reduction in peak torque). Both the *unfatigued* and *fatigued runs* were performed in a *traditional* shoe and in *advanced footwear technology* on an instrumented treadmill while three-dimensional motion capture and ground reaction force data were collected.

**Results:** Plantar flexor fatigue led to a redistribution of joint work, with lower positive ankle work (p < 0.001), greater positive knee work (p = 0.001), and similar positive hip work (p = 0. 550) in the *fatigued run* as compared to the *unfatigued run*. The relative positive joint contributions shifted proximally with fatigue (ankle - 4%, knee +3%); however, this shift was not influenced by shoe condition (p > 0.05).

**Conclusion:** Plantar flexor fatigue contributes to the proximal redistribution of positive joint work during running. However, the adoption of a suboptimal gait strategy appears to occur regardless of whether a runner wears a traditional shoe or advanced running footwear technology.

## Introduction

Mechanical work is an important biomechanical determinant of running performance. For example, it explains ∼76% of the metabolic cost of running at a range of submaximal velocities (2.2-4.6 m s^−1^ (1)). In unfatigued running (3.5 m s^−1^), the ankle joint generates approximately 54% of the positive work required, while the knee and hip joint each generate approximately 23% (2). Mechanical work from the lower limb joints is likely organized in a manner that promotes metabolically efficient running. The ankle joint, for example, is well suited to its principal role in positive power generation. The Achilles tendon enhances running efficiency by storing and releasing elastic energy (3–6), which lowers demands on muscles and allows for substantial energy savings (6–9). Moreover, the plantar flexor muscles have shorter fascicles and lower volumes than more proximal muscles (10), leading to a lower metabolic cost of contraction (11). For these reasons, the muscle-tendon units of the knee and hip are considered to be less metabolically efficient (11).

As an exhaustive run progresses, positive joint work shifts proximally from the ankle to the knee—a movement strategy that is likely more metabolically costly. This redistribution has been observed in recreational and trained male and female runners, over various distances (e.g., 3-10 km) (12–14). The latter stages of exhaustive running are also associated with higher metabolic cost relative to the earlier stages (15–19), suggesting that the shift in joint work may contribute to this energetic penalty. However, establishing a causal relationship remains challenging. Through its potential influence on metabolic cost, the proximal shift in joint work is thought to reduce running performance (12–14, 20).

Neuromuscular fatigue (i.e., a reduction in force or power output from a muscle group (21)) of the plantar flexors is purported to trigger the proximal redistribution of positive joint work in running (12–14, 20). In support of this hypothesis, a proximal shift in positive joint work is observed at higher intensity running bouts corresponding to 10 km race pace or faster (12–14, 20), but not after a prolonged (20 km) run at a self-selected ‘easy’ pace (2). Whether plantar flexor fatigue is indeed responsible for the redistribution of positive joint work during running remains unknown.

Advanced footwear technology (AFT) may affect the positive redistribution of ankle work caused by plantar flexor fatigue. Indeed, the proximal redistribution of positive joint work during a fatiguing run is delayed (but not prevented) when running in a shoe with a carbon fiber plate placed beneath the insole (14). Compared to traditional running shoes, AFT reduces negative and positive ankle work, yet maintains the relative percentage joint contribution (22, 23). A runner’s ability to retain a favorable distribution of mechanical work under conditions of local muscle fatigue could help explain the substantial impact of AFT on performance (24, 25).

The first aim of this study was to determine the effect of bilateral plantar flexor fatigue on lower limb positive joint work during running. We hypothesized that, compared to unfatigued running, positive ankle joint work would decrease and positive knee joint work would increase (12–14, 20) under conditions of plantar flexor fatigue. We sought to induce local plantar flexor fatigue at levels commensurate to that accumulated during a 5-10 km run (19-27% fatigue; (26–28)). Furthermore, given that work redistribution occurs primarily when running at high intensities (2, 12, 13), we focused on shorter, faster runs.

Acknowledging the effect of AFT on running performance (24, 25), metabolic cost (29–31), and biomechanics (22, 23), the second aim of this study was to determine the effect of *traditional* shoes versus *AFT* on lower limb positive joint work during running with plantar flexor fatigue. We hypothesized that compared to a *traditional* shoe, the anticipated decrease in positive ankle work and increase in positive knee work would be less pronounced in *AFT* when running with local plantar flexor fatigue. While positive work contributes significantly to the metabolic cost of running (32), negative work also contributes but to a lesser extent (33, 34). Given that both are influenced by AFT (22), a tertiary analysis of negative ankle, knee and hip joint work was conducted.

A greater understanding of the mechanisms that underpin the deterioration of gait mechanics during running may inform the development of preventative techniques. Identifying the potential benefits of *AFT* on these determinants of running performance may lead to further footwear innovations.

## Methods

### Participants

All study procedures were approved by the University of New South Wales Human Research Ethics Committee (iRECS6464). Healthy male participants (*n* = 12; age: 26.0 ± 4.4 years; body mass: 77.2 ± 4.7 kg; height: 1.84 ± 0.04 m; mean ± sd; see Supplementary Table 1) provided written informed consent after receiving clear explanations of the study procedures. An a priori power analysis (G*Power) with an alpha level of 0.05, power of 0.80, and an effect size of 0.88 was conducted for positive ankle work (12). A sample size of 13 participants was determined, consistent with previous research that found a proximal shift in positive joint work during the later stages of a 10 km exhaustive run (12).

Participants were required to have a 10 km race time between 35 and 45 min and be free of any respiratory, cardiovascular, neurological, or other conditions that could affect gait. Additional inclusion criteria included a shoe size between US8 and 12, no lower limb injuries in the past two months, and no history of rhabdomyolysis. Females have lower muscle fatiguability than males (35–37), and fatiguability is affected by the specific phase of the menstrual cycle (38), which is often highly individual (39). Because this study was initially planned as part of larger multiday study, which would be affected by day-to-day variations, only males were recruited.

### Participant preparation

Participants were instructed to refrain from fatiguing exercise the day before and the day of testing. Anthropometric measurements, such as body mass and height were recorded, and limb dominance was determined (40). Participants were fit for both the *traditional* shoe (Brooks Hyperion Tempo^TM^) and *AFT* (ASICS METASPEED SKY+^TM^). Shoes were matched for mass and the order they were tested in was randomly assigned to each participant. Shoe characteristics are shown in Figure 1 (41–44).

**Figure 1.**
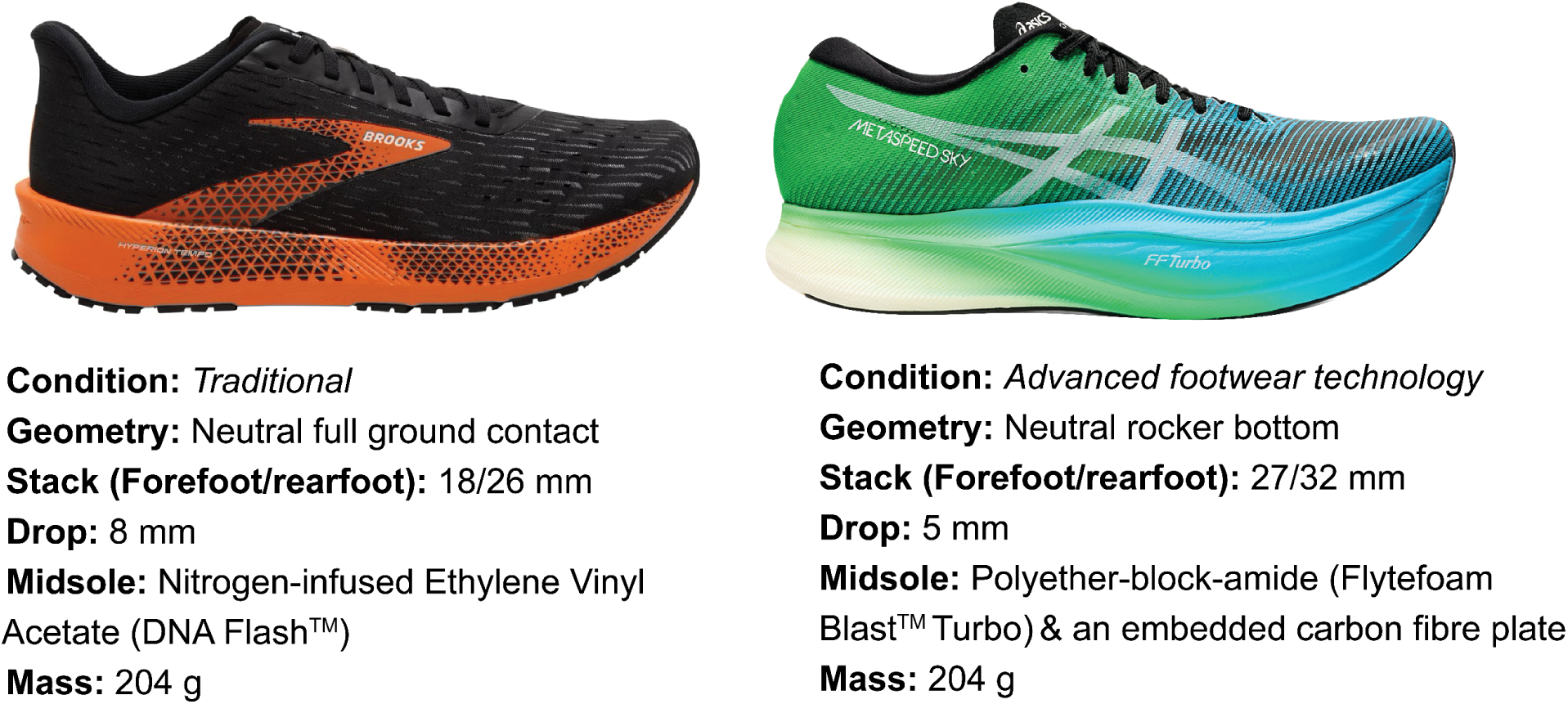
Traditional shoe (Brooks Hyperion Tempo^TM^; left) and advanced footwear technology (AFT; ASICS METASPEED SKY+^TM^; right) characteristics. Stack = the distance from outsole to the insole, midsole = material(s) between the outsole and insole, drop = height of the rearfoot stack minus height of the forefoot stack (i.e., shoe gradient), mass = mass of a single men’s US9 shoe. Stack, drop and mass based on measurements from Running Warehouse (41–44).

Retro-reflective markers were placed over the C7 and T10 spinous process, the clavicular notch and the xiphoid process. They were also placed bilaterally over the acromion process, the anterior superior iliac spine, the posterior superior iliac spine, the iliac crest, the thigh (3 marker cluster), the medial and lateral femoral epicondyle, the shank (3 marker cluster), the medial and lateral malleoli, the medial, lateral and posterior calcaneus, the first metatarsal phalangeal joint, the fifth metatarsal phalangeal joint and the tip of the hallux (12, 20, 45). The foot markers were attached to the corresponding positions on the shoe.

Participants completed a 5 min warm-up run on the treadmill (M-Gait, Motek Medical, DIH, Amsterdam, Netherlands) at self-selected speeds between 10 and 12 km h^−1^ (2.77 and 3.33 m s^−1^) with a 0° incline. For safety purposes, they were also familiarized with the belt acceleration and deceleration during a series of six brief (∼30 s) trials. Following this they were familiarized with the position they were to adopt in the dynamometer, and with the MVC protocol (described below).

### Experimental set-up

Marker trajectories were tracked in three-dimensions using an eight-camera infrared motion capture system (200 Hz; Bonita, Vicon Motion Systems, Oxford, UK). Synchronized ground reaction force (GRF) data were collected via an instrumented split-belt treadmill (1000 Hz; M-Gait, Motekforce Link, DIH, Amsterdam, Netherlands).

Peak plantar flexor torques during maximum voluntary contractions (*MVC*s) were measured via a custom dynamometer (1000 Hz). Each MVC was performed for 3-5 s with verbal encouragement provided throughout the contractions (46). Trials were separated by a 1 min rest period (47, 48). Participants were seated with their hips flexed at 90°, knees maximally extended, talocrural joint positioned at 0°, and their foot secured to a footplate against which they plantarflexed. The dynamometer setup was adjusted to fit individual anthropometric measurements. A seat belt was fastened around the participants’ waist to limit sliding in the seat. The dynamometer was positioned next to the treadmill and fatigue protocol equipment (i.e., step), enabling a rapid transition of approximately 10 s between activities to minimize the decay of fatigue (49).

### Experimental protocol

The protocol is outlined in Figure 2. Three *MVCs* were performed in the first shoe, and then three in second shoe (order randomized); this was to account for the different properties (e.g., compliance, stack height) of the shoes that could potentially affect maximum torque production.

**Figure 2.**
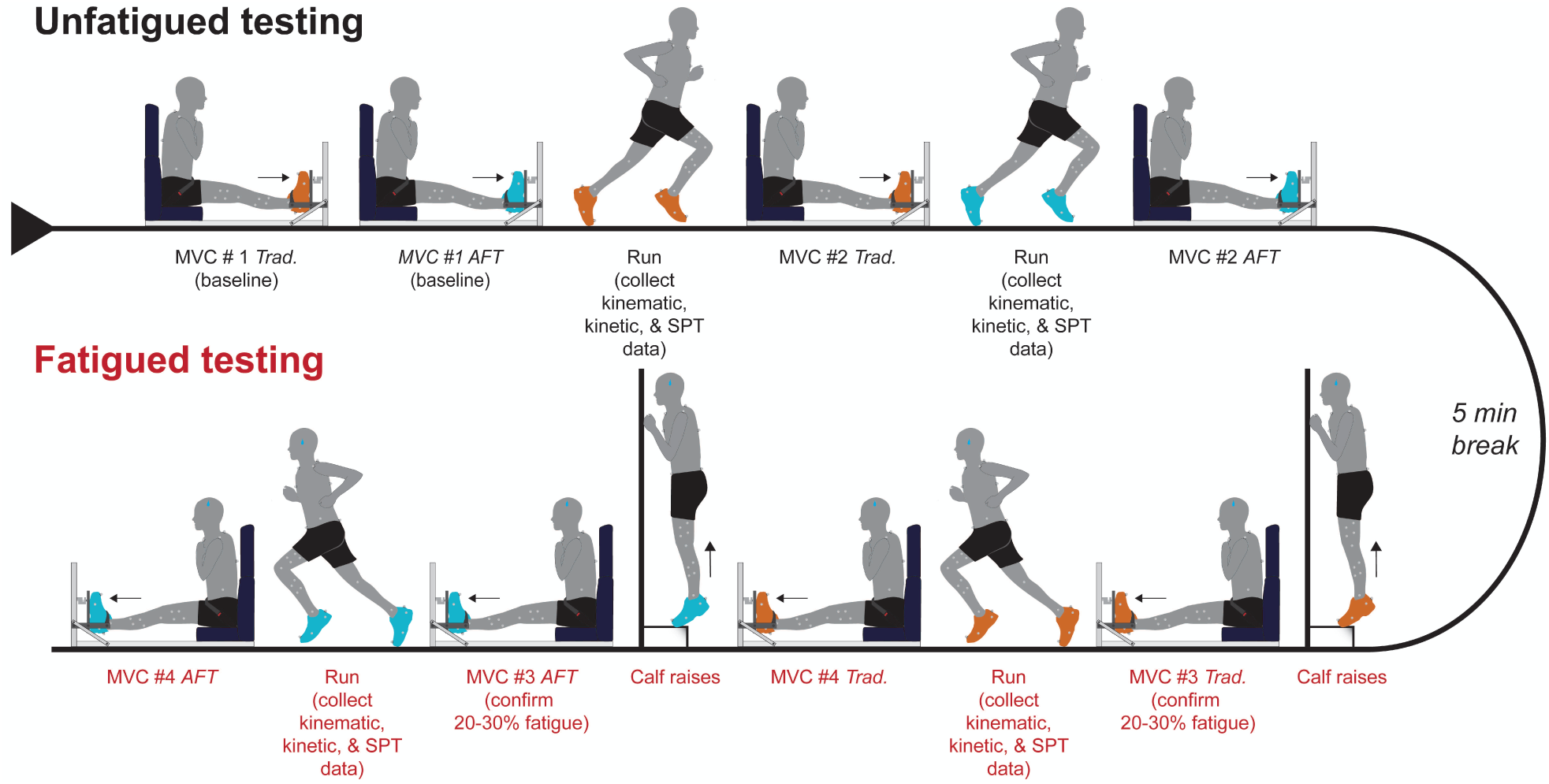
Study protocol conducted over a single day. Kinematic, kinetic and spatiotemporal data during unfatigued and fatigued runs in the traditional (trad.) shoe (orange) and advanced footwear technology (AFT; blue). Shoe order was randomised with the traditional shoe displayed first in the above figure. The fatigue protocol consisted of repeated calf raises to failure. MVC = maximum voluntary contraction.

The highest torque value achieved in any contraction for each shoe was recorded as the baseline peak torque measurement for the corresponding shoe (*MVC #1*).

After baseline *MVC* testing, participants rested for 5 min. Next, they completed static calibration trials and 30 s *unfatigued runs* at their individual 10 km race pace (mean: 4.0 m s^−1^, range: 3.7-4.5 m s^−1^). A static calibration and run were conducted in each shoe, in the same order as the baseline *MVCs* (AB or BA). Immediately following each run, participants performed a *MVC* (*MVC #2*). For five participants, the highest recorded torque value, in at least one shoe, occurred during *MVC #2*, which was subsequently used as the baseline measure. Shoes were changed during a 5 min rest between the post-*unfatigued run MVC* in the first shoe and the *unfatigued run* in the second shoe.

Next, participants underwent a fatigue protocol consisting of two sets of bilateral bodyweight calf raises on the edge of a 20 cm step while wearing the first shoe (50). For balance, they were permitted to lightly touch the wall in front of them (Fig. 2). Repetitions were performed to a 60-bpm metronome, with beats timed to full plantarflexion and dorsiflexion (51). Verbal encouragement was provided. Each set was continued until failure, defined either volitionally or by the researcher (e.g., incomplete range of motion or inability to maintain timing with the metronome). After completing the two sets, with 1 min rest between sets, participants immediately performed an *MVC* (*MVC #3*) to confirm that plantar flexor torque output had decreased to 70-80% of the peak baseline measurement. This range was selected to reflect typical torque reductions after a near-maximal 10 km run (26–28). If the *MVC* measure was not in the fatigue range, participants either completed additional sets or rested briefly before repeating the *MVC* assessment. Once the required fatigue range was achieved, participants immediately transitioned to the *fatigued run* on the treadmill. The conditions for the *fatigued run* mirrored those of the *unfatigued run*.

After the *fatigued run* in the first shoe, participants performed a final *MVC* (*MVC #4*). They then switched to the second shoe and repeated the fatigue protocol. If their fatigue level remained within the target range, they proceeded with the *fatigued run* in the second shoe. Otherwise, an additional set of calf raises and *MVC* assessments were performed until the suitable fatigue range was achieved. Immediately after, they completed the *fatigued run* in the second shoe, the final *MVC* was recorded.

During both *unfatigued* and *fatigued runs,* marker trajectory and GRF data were captured.

### Data analysis

During dynamometer testing, force (N) was measured and converted to torque (Nm). The moment arm was defined in the static calibration trials for each shoe condition. To obtain the moment arm, the distance between the medial malleolus and first metatarsophalangeal markers formed the hypotenuse. The horizontal component, adjacent to the angle between the hypotenuse and the horizontal axis, was taken as the effective moment arm.

Fatigue was defined as a decline in torque relative to the torque baseline measurement taken from the highest peak measurement across *MVC #1* or *#2* of the corresponding shoe. For the *traditional* shoe, eight of twelve participants had their baseline peak torque measurement recorded in *MVC #1* and four participants had it recorded in *MVC #2.* For the *AFT*, nine of twelve participants had their baseline peak torque measure recorded in *MVC #1* and three had it recorded in *MVC #2*. Percentage changes in peak plantar flexor torque were calculated using the raw data.

All marker trajectories and GRF data were filtered using a dual pass fourth-order Butterworth low-pass filter with a 20 Hz cutoff frequency. Data were analyzed over the last 20 stance phases of each run. Spatiotemporal variables, including stride length (m), stride time (s), contact time (s), and flight time (s) were computed from force plate-derived events (stride/contact/flight time) and force plate-derived events in conjunction with marker data (stride length). Joint angles and net moments, total power and work were calculated via custom MATLAB (R2022b, MathWorks, Natick, MA, USA) scripts that included inverse dynamics analysis (12, 45). For this, segment inertial parameters were defined from each body according to body mass and height (52). Joint centres were defined using regression-based methods for the hip (53) and anatomical landmarks for the knee and ankle. Joint moments were expressed in the proximal segment’s anatomical coordinate system, and sagittal-plane GRF moment arm lengths were defined relative to this system. Moment arm lengths were calculated by dividing the GRF term from Hof’s explicit joint torque equation (54) by the GRF vector magnitude, yielding the perpendicular distance from the joint centre to the GRF line of action.

To avoid overestimating ankle joint power, the ankle joint complex approach was used, i.e., shank-calcaneus power (55). Joint angles were referenced to a neutral position determined from a standing posture during a static calibration trial, with separate static trials performed for each shoe, and joint moments were expressed in the anatomical coordinate system of the proximal segment. Positive and negative joint work at the ankle, knee, and hip were calculated via numerical integration of the total ankle joint power-time curve, with positive work representing the sum of all positive integrals and negative work the sum of all negative integrals.

Furthermore, relative positive ankle, knee and hip joint work was computed as the percent contribution of joint work relative to the summed ankle, knee and hip joint work.

Stride lengths were non-dimensionalized by *L* and time variables were non-dimensionalized by 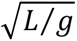. Power data were non-dimensionalized by 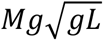, while work and moments were non-dimensionalized by *MgL*, where *M* is body mass, *g* is acceleration due to gravity, and *L* is leg length (distance between the medial malleolus and anterior superior iliac spine markers of the dominant limb). Non-dimensionalized work outputs were further normalized by non-dimensionalized stride length (i.e., *stride lenght/L*) to account for potential variations in distance traveled between runs.

### Statistical analysis

All statistical analyses were performed using linear mixed effects models in the R programming language (R Development Core Team, Vienna, Austria). Specifically, the lmer function from the lme4 package and the emmeans function from the emmeans package were used (56, 57). For each model, participants were modelled as a random factor with a random intercept. The threshold for statistical significance was set to p < 0.05. Normality was evaluated and confirmed via inspection of residual and Q-Q plots. Group data are presented as estimated marginal means [95%CI] of the predicted values from the linear mixed effects model.

Categorical factors in the models included run (*unfatigued* or *fatigued*) and shoe (*traditional* or *AFT*). Seven dependent variables were analyzed: positive and negative joint work (ankle, knee, and hip), peak plantar flexor torque, stride length, stride time, contact time and flight time. Each model assessed the fixed effects of the run condition, shoe type, and their interaction effect on the individual dependent variables.

A likelihood ratio test was conducted to assess model fit with and without a run and shoe interaction. If model fit was improved by the inclusion of the interaction (p < 0.05), it was retained and post-hoc comparisons were assessed for each shoe, between runs. If model fit was not improved, the interaction was removed, data was pooled across shoes and post-hoc comparisons were assessed between runs. Custom contrasts were specified for peak plantar flexor torque (i.e., *MVC* #*2*–*1*, *MVC* #*4*–*3, MVC* #*4*–*2, MVC #1 Shoe B -MVC #1 Shoe A*) to constrain the model and investigate relevant comparisons. Bonferroni correction was applied.

## Results

### Maximum plantar flexor torque

The estimated marginal means for peak plantar flexor torque are presented in Table 1. The interaction between run and shoe did not improve model fit (p = 0.928). Baseline peak plantar flexor torque (MVC #1) was similar between shoes (p = 0.065), with a mean difference of −2 Nm [−5.1, 0.2]. Thus, shoe condition had no effect on baseline torque.

**Table 1.** Estimated marginal means for peak plantar flexor torque (N m [95%CI]) for *maximum voluntary contractions (MVCs*) #*1-4* for the *pooled* shoe data*, traditional (Trad.)* shoe and *advanced footwear technology (AFT)* for the cohort (*n =* 12). Differences (mean [95%CI]) and associated p-values are provided for *MVC #2*-*1*, *4*-*3, and #4-2* for the *pooled* group as an interaction between shoe and *MVC #* was removed.

| Group | *MVC #1* | *MVC #2* | *MVC #3* | *MVC #4* | Difference<br>(*MVC #2-1*) | Difference<br>(*MVC #4-3*) | Difference<br>(*MVC #4-2*) |
| --- | --- | --- | --- | --- | --- | --- | --- |
|  | (N m) |  |  |  |  |  |  |
| *Pooled* | 86<br>[71, 101] | 85<br>[70, 99] | 66<br>[51, 81] | 72<br>[58, 87] | -1<br>[-5, 3] | 6*<br>[2, 10] | -12*<br>[-17, -8] |
| *Trad.* | 87<br>[72, 102] | 86<br>[71, 101] | 68<br>[53, 83] | 74<br>[59, 89] |  |  |  |
| *AFT* | 85<br>[70, 99] | 84<br>[68, 98] | 65<br>[50, 80] | 71<br>[56, 86] |  |  |  |

\*Indicates statistical significance ( $p < 0.05$ ).

Peak plantar flexor torque was similar when measured before (*MVC #1*) and after (*MVC #2*) the *unfatigued run* (p = 0.524), indicating no accumulation of plantar flexor fatigue during the *unfatigued run*. The fatigue protocol generated 24% [22, 26] plantar flexor fatigue for both shoes. Furthermore, peak plantar flexor torque was lower by 7% [5, 9] when measured before (*MVC #3*) as compared to after the *fatigued run* (*MVC #4*; p = 0.003). However, it was higher by 13% [11, 14] when measured after the *unfatigued run* as compared to after the *fatigue run* (p < 0.001). Therefore, plantar flexor fatigue recovered over the *fatigued run* but not to unfatigued levels.

### Joint mechanics

All parameters except joint angles are non-dimensionalized (see Data analysis section) and thus are unitless. Figure 3 presents group mean joint angles, moments and powers at the ankle, knee and hip for the two shoe conditions (*traditional, AFT*) during the stance phase of the *unfatigued* and *fatigued runs*.

**Figure 3.**
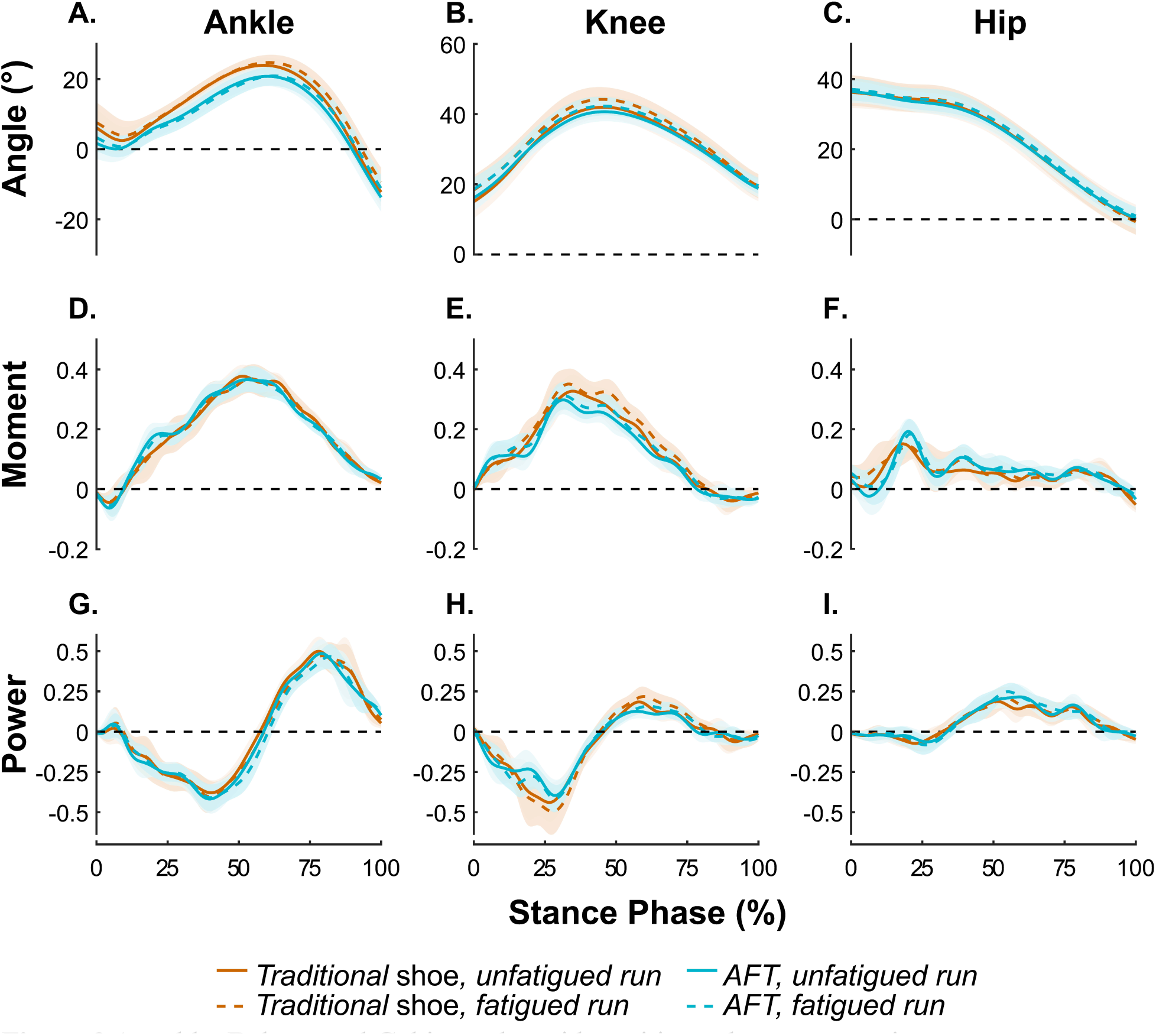
**A.** ankle, **B.** knee and **C**. hip angles with positive values representing flexion/dorsiflexion and **D.** ankle, **E.** knee and **F.** hip net moments (non-dimensionalized) with positive values representing extensor/plantar flexor moments. **G.** Ankle, **H.** knee and **I.** hip total power (non-dimensionalized) with positive values representing power generation. Mean [95%CI] (shaded regions) data are presented for (*n* = 12). The unfatigued (solid line) and fatigued (dashed lines) run for the traditional shoe (orange) and AFT (blue) conditions are shown over the stance phase (foot strike to foot off).

### Lower limb joint positive joint work

The estimated marginal means for positive ankle, knee and hip work are reported in Table 2. The interaction between run and shoe did not improve model fit for positive ankle (p = 0.817), knee (p = 0.253), and hip (p = 0.845) work. This indicates that shoe condition did not influence positive joint work distribution when running without and with plantar flexor fatigue. When pooled across the shoes, positive ankle work was 4% lower (p < 0.001), knee work was 3% higher (p = 0.001) and hip work was similar (p = 0.550) in the *fatigued run* as compared to the *unfatigued run*.

**Table 2.** Estimated marginal means [95%CI] for positive ankle, knee and hip work (non-dimensionlized) during stance phase of the *unfatigued* and *fatigued run* for *pooled* shoe data*, traditional* shoe *(Trad.)* and *advanced footwear technology (AFT)* for the cohort (*n =* 12). Differences (mean [95%CI]) between *unfatigued* (*UFR*) and *fatigued (FR) runs* are provided for the *pooled* shoe data as an interaction between shoe and run condition was removed.

| Shoe | Run | Positive ankle<br>work | Difference<br>(*FR-UFR*) | Positive knee<br>work | Difference<br>(*FR-UFR*) | Positive hip<br>work | Difference<br>(*FR-UFR*) |
| --- | --- | --- | --- | --- | --- | --- | --- |
| (Unitless; non-dimensionalized) |  |  |  |  |  |  |  |
| *Pooled* | *UFR* | 0.0310 | -0.0024* | 0.0099 | 0.0020* | 0.0192 | 0.0007 |
|  |  | [0.0284, 0.0336] | [-0.0036, -0.0012] | [0.0080, 0.0119] | [0.0089, 0.0031] | [0.0167, 0.0218] | [-0.0016, 0.0030] |
|  | *FR* | 0.0287 |  | 0.0119 |  | 0.0199 |  |
|  |  | [0.0261, 0.0313] |  | [0.0010, 0.0138] |  | [0.0174, 0.0225] |  |
| *Trad.* | *UFR* | 0.0323 |  | 0.0114 |  | 0.0186 |  |
|  |  | [0.0297, 0.0350] |  | [0.0094, 0.0134] |  | [0.0159, 0.0214] |  |
|  | *FR* | 0.0300 |  | 0.0134 |  | 0.0193 |  |
|  |  | [0.0273, 0.0350] |  | [0.0114, 0.0154] |  | [0.0166, 0.0221] |  |
| *AFT* | *UFR* | 0.0297 |  | 0.0085 |  | 0.0198 |  |
|  |  | [0.0271, 0.0324] |  | [0.0065, 0.0105] |  | [0.0177, 0.0232] |  |
|  | *FR* | 0.0274 |  | 0.0105 |  | 0.0205 |  |
|  |  | [0.0247, 0.0300] |  | [0.0085, 0.0124] |  | [0.0177, 0.0232] |  |

\*Indicates statistical significance ( $p < 0.05$ ).

### Lower limb joint negative joint work

The estimated marginal means for negative ankle, knee and hip work are listed in Table 3. The interaction between run and shoe did not improve model fit for negative ankle (p = 0.837), knee (p = 0.733), and hip (p = 0.539) work. Once again, this indicates that shoe condition did not influence negative joint work distribution when running without and with plantar flexor fatigue. When pooled across the shoes, negative ankle work was similar between runs for the ankle (p = 0.682), knee (p = 0.307), and hip (p = 0.443) in the *fatigued run* compared to the *unfatigued run*.

**Table 3.**
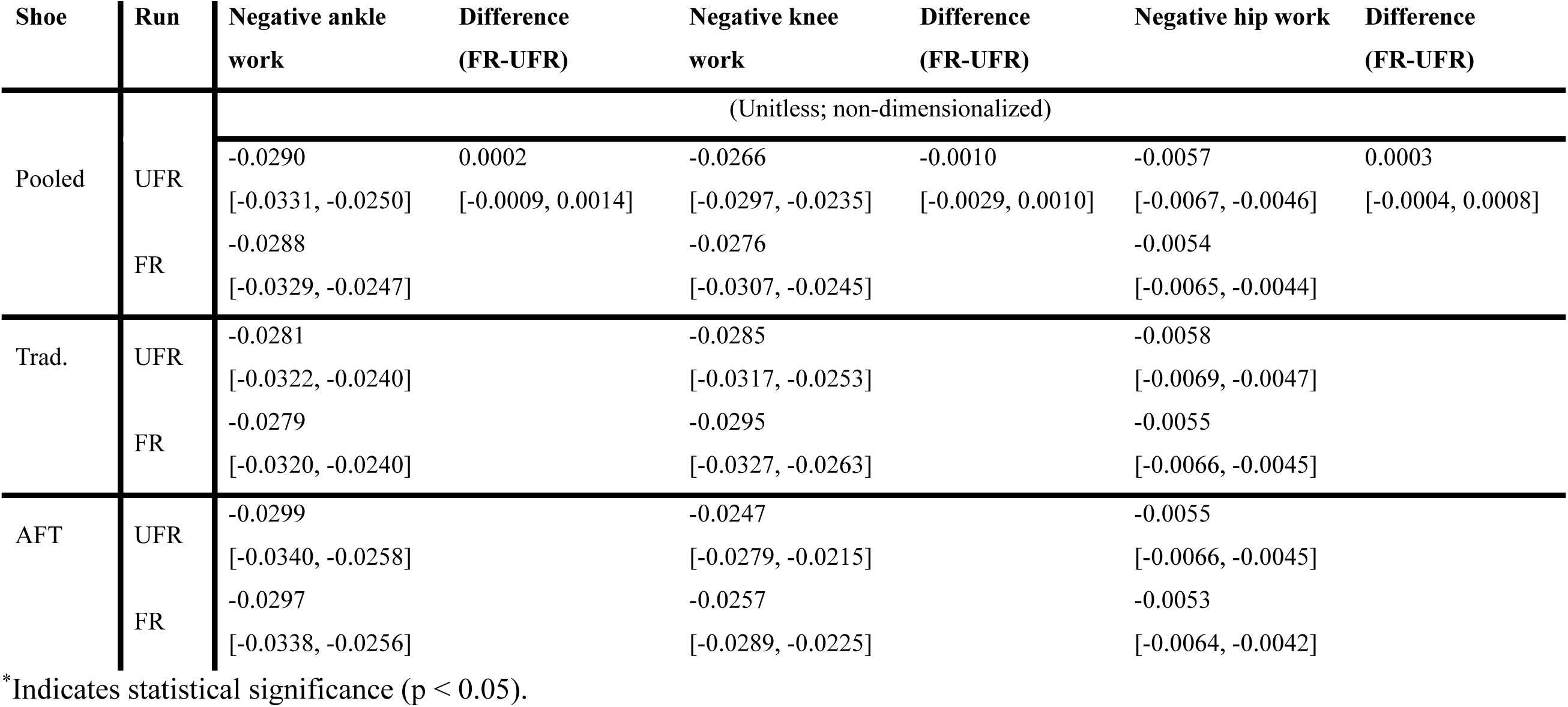
Estimated marginal means [95%CI] for negative ankle, knee and hip work (non-dimensionlized) during stance phase of the *unfatigued* and *fatigued run* for *pooled* shoe data*, traditional* shoe (*Trad*.) and *advanced footwear technology (AFT)* for the cohort (*n = 12*). Differences (mean [95%CI]) between the *unfatigued (UFR)* and *fatigued (FR) runs* are provided for the *pooled* shoe data as an interaction between shoe and run condition was removed.

### Spatiotemporal variables

The interaction between run and shoe did not improve model fit for stride length (p = 0.631), stride time (p = 0.483), contact time (p = 0.937), and flight time (p = 0.381). Therefore, shoe condition did not influence spatiotemporal variables during running without and with plantar flexor fatigue.

As seen in Table 4, when pooled across the shoes, stride length (p = 0.899), stride time (p = 0.620), contact time (p = 0.073), and flight time (p = 0.431) were similar in the *fatigued run* and the *unfatigued run*. Therefore, plantar flexor fatigue did not affect spatiotemporal variables.

**Table 4.** Estimated marginal means [95%CI] for stride length, stride time, contact time and flight time (non-dimensionlized) during the *unfatigued* and *fatigued run* for *pooled* shoe data*, traditional* shoe (*Trad*.) and *advanced footwear technology (AFT)* for the cohort (*n =* 12). Differences (mean [95%CI]) between the *unfatigued (UFR)* and *fatigued (FR) runs* are provided for the *pooled* shoe data as an interaction between shoe and run condition was removed. The estimated marginal means for flight time appear identical across conditions, which is attributable to minimal effect sizes and rounding.

| Shoe | Run | Stride time | Difference<br>(FR-UFR) | Stride length | Difference<br>(FR-UFR) | Contact time | Difference<br>(FR-UFR) | Flight time | Difference<br>(FR-UFR) |
| --- | --- | --- | --- | --- | --- | --- | --- | --- | --- |
| (Unitless; non-dimensionalized) |  |  |  |  |  |  |  |  |  |
| *Pooled* | *UFR* | 2.19 | -0.01 | 2.90 | 0.00 | 0.644 | -0.013 | 1.55 | 0.01 |
|  |  | [2.12, 2.26] | [-0.03, 0.02] | [2.79, 3.01] | [-0.04, 0.04] | [0.605, 0.684] | [-0.027, 0.001] | [1.49, 1.61] | [-0.01, 0.02] |
|  | *FR* | 2.18 |  | 2.89 |  | 0.631 |  | 1.55 |  |
|  |  | [2.12, 2.25] |  | [2.78, 3.00] |  | [0.592, 0.671] |  | [1.49, 1.61] |  |
| *Trad.* | *UFR* | 2.20 |  | 2.91 |  | 0.651 |  | 1.55 |  |
|  |  | [2.13, 2.27] |  | [2.79, 3.01] |  | [0.611, 0.692] |  | [1.49, 1.61] |  |
|  | *FR* | 2.19 |  | 2.90 |  | 0.638 |  | 1.55 |  |
|  |  | [2.12, 2.26] |  | [2.79, 3.01] |  | [0.598, 0.679] |  | [1.49, 1.61] |  |
| *AFT* | *UFR* | 2.18 |  | 2.89 |  | 0.638 |  | 1.55 |  |
|  |  | [2.12, 2.25] |  | [2.78, 3.00] |  | [0.597, 0.678] |  | [1.49, 1.61] |  |
|  | *FR* | 2.18 |  | 2.89 |  | 0.625 |  | 1.55 |  |
|  |  | [2.11, 2.25] |  | [2.77, 3.00] |  | [0.584, 0.665] |  | [1.49, 1.61] |  |

315 \*Indicates statistical significance ( $p < 0.05$ ).

## Discussion

Plantar flexor fatigue that accumulates during a prolonged run at a near-maximal effort (e.g., 10 km race) has been proposed to cause a proximal redistribution of positive joint work (12–14, 20). This shift in positive joint work from the ankle to the knee, may inflate metabolic cost (32, 58) and reduce performance (12–14, 20). The first aim of the current study was therefore to quantify the effect of bilateral plantar flexor fatigue on lower limb positive joint work during running. We successfully induced plantar flexor fatigue (∼24%) comparable with that observed after exhaustive bouts of running (26–28). In line with our hypothesis, in a cohort of trained male runners, this fatigue caused a decrease in positive ankle work, an increase in positive knee work and no change in positive hip work (Table 2). Relative contributions to total positive work decreased by 4% at the ankle and increased by 3% at the knee (Fig.4)

**Figure 4.**
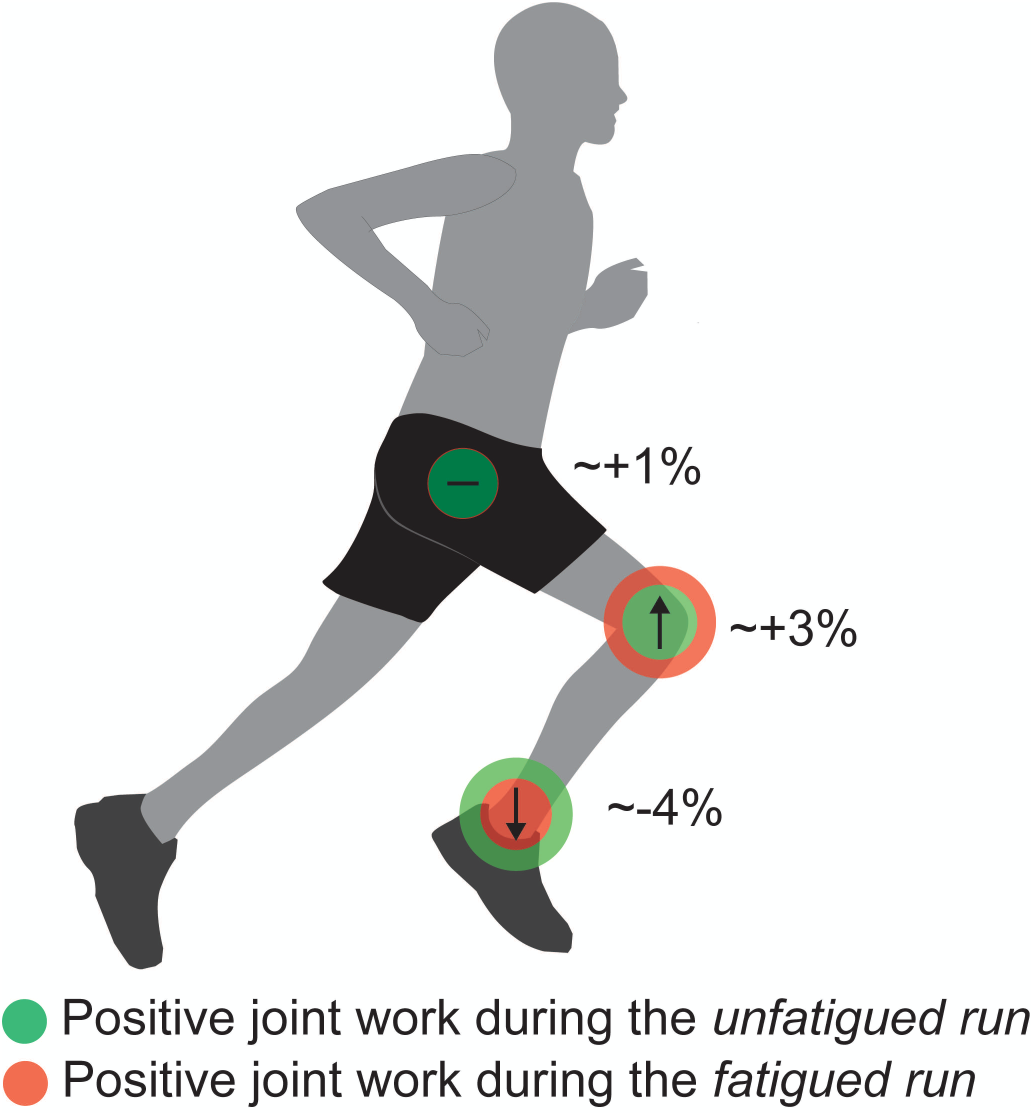
Relative positive joint work redistribution at the ankle, knee, and hip for *Pooled* shoe data over the *fatigued run* and *unfatigued run*.

The results are consistent with findings from prior studies that used running-induced global fatigue protocols (e.g., 10 km exhaustive run) in similarly trained runners (12, 14). Although fatigue was not directly measured in these studies, the level of plantar flexor fatigue induced by similar running protocols (12, 14) (i.e., 19-27% reduction in peak plantar flexor torque, (26–28)) is comparable with the current study (24%). Thus, plantar flexor fatigue, whether present in conjunction with fatigue in other muscle groups (global) or present in isolation (local), is associated with a proximal shift in positive joint work during running.

The second aim was to determine the effect of a *traditional* shoe versus *AFT* on lower limb positive joint work during a run with plantar flexor fatigue. To our surprise, the shift in positive joint work when running with plantar flexor fatigue was similar between the shoes. Finally, we observed no differences in negative joint work when running without versus with plantar flexor fatigue.

Neuromuscular fatigue has long been recognized as a determinant of running performance (59) and the current study offers new mechanistic insight. Specifically, we provide evidence that plantar flexor fatigue contributes to the proximal redistribution of positive joint work occurring during an exhaustive run, a shift that likely undermines running efficiency. Joint work explains the majority of the metabolic cost of running (1), and a more distal distribution is likely associated with lower energy demands (4–6, 11). Hence, this shift could have meaningful performance implications, especially during the final stages of a race. Recently, a potential fourth physiological determinant of performance has been identified: metabolic durability, which refers to the ability to resist increases in metabolic cost over the course of prolonged exercise (60). Our results may have important implications for enhancing this determinant and understanding the mechanisms by which strength training (e.g., calf raises) improves it (61). We recently found no effect of plantar flexor fatigue on running economy at slower speeds (∼10–12 km·h⁻¹; Nahan et al., unpublished). However, the higher velocities in the current study (∼14.4 km·h⁻¹) likely imposed greater mechanical demands and amplified the effects of plantar flexor fatigue (13). Indeed, positive joint work redistribution primarily occurs when running at a near-maximal effort (12–14, 20), not slower, easy efforts (2). Although the precise energetic cost of this shift remains unknown due to the limitations of indirect calorimetry in capturing anaerobic metabolism, these gait adaptations are likely metabolically suboptimal.

The anticipated protective effect of AFT on the fatigue-induced redistribution of relative joint work was not observed. Thus, running shoes may have limited influence on this gait parameter once substantial plantar flexor fatigue has developed. While *AFT* features, such as the embed stiffening agent (associated with increased longitudinal bending stiffness), may delay the onset of plantar flexor fatigue during running (14), they do not appear to prevent it. It is currently unclear whether a 10 km exhaustive run in AFT results in similar levels of plantar flexor fatigue as compared to running in traditional shoes. Hence, the performance benefits observed with AFT may be linked to a preservation of positive joint work distribution, potentially due to reduced plantar flexor fatigue. Future research should explore this further. In our study, fatigue was induced directly, which eliminated the possibility for *AFT* to delay the onset of plantar flexor fatigue and the associated proximal shift in positive joint work. However, during a fatiguing run, this delay (14) may help explain the reported metabolic (29–31) and performance (24, 25) benefits of *AFT*. Interestingly, greater plantar flexor fatigue occurs over a half marathon in shoes with a carbon fiber plate as compared to those without a plate (62), however, these were flat, isolated plates placed in traditional shoes (14, 62), rather than the fully integrated AFT design (i.e., a curved embedded stiffening agent in a novel midsole foam) design used here. Our findings indicate that in the final stages of a race, once substantial plantar flexor fatigue has accumulated, shoe choice does not meaningfully influence the relative distribution of positive joint work and the negative impact it may have on running performance.

### Limitations

The foot can contribute up to 2.6% of total positive power generation in running (14) and changes in foot work may occur with fatigue (e.g., at the metatarsophalangeal joint (14)). However, in line with many previous studies exploring the redistribution of mechanical work during running (2, 12, 13, 20), we did not account for foot power contributions due to the technical complexity. Also, the effect of our local fatigue protocol (calf raises) on the mechanical contributions of the foot during a run is unclear.

Cyclic stress and strain on musculotendon units from the stretch-shortening cycle results in complex neuromuscular fatigue accumulation, which may differ from fatigue induced by calf raises (63). However, replicating running-specific stresses with a single-joint task is not feasible, nor was it the aim of this study.

The plantar flexors recovered by 7% over the course of the *fatigued run*, which may partially explain why previous research on running-induced fatigue found greater distal-to-proximal positive joint work shifts than in the current study (12–14, 20). Nevertheless, 17% plantar flexor fatigue measured post-*fatigued run* was substantial and likely sufficient to meaningfully impact running gait. Furthermore, plantar flexor torque did not recover to unfatigued levels.

Isometric MVCs are commonly used to assess neuromuscular fatigue (64); however, some argue that dynamic measures, such as concentric or eccentric plantar flexor contractions in an isokinetic dynamometer, may better capture local torque changes following running (65, 66). Yet, the plantar flexors contract in a quasi-isometric manor during running (67, 68) and previous studies measuring plantar flexor fatigue after 10 km exhaustive runs have utilized isometric MVCs (26–28). We adopted the same protocol to be consistency with these studies.

Since the menstrual cycle is known to influence neuromuscular fatigue (38, 39), the present study focused solely on male runners. As such, the present findings cannot be directly extrapolated to female runners and future research should address this gap in the literature.

## Conclusion

This study is the first to empirically investigate the hypothesis that plantar flexor fatigue is key to the proximal redistribution of positive joint work observed during exhaustive running. We found that, when running with bilateral localized plantar flexor fatigue, positive ankle joint work decreased and positive knee joint work increased, yet the shift in joint work was not affected by wearing a *traditional* running shoe or *AFT.* Our findings offer deeper insight into the mechanisms underlying performance impairments during exhaustive running bouts. This underscores the importance of plantar flexor endurance training (e.g., calf raises) and suggest that running shoes may not be sufficient to offset high magnitudes of plantar flexor fatigue. Future research could consider the extent to which changes in positive joint work distribution influence running performance, and what magnitude of change is necessary to produce a meaningful deterioration in performance.

## Acknowledgements

We acknowledge Hiu Ying Or for aiding in data collection, as well as the Morcom Duncan Family Scholarship for partially funding the research.

## References

1. Riddick RC, Kuo AD. Mechanical work accounts for most of the energetic cost in human running. Sci Rep. 2022;12(1):645.

2. Melaro JA, Gruber AH, Paquette MR. Joint work is not shifted proximally after a long run in rearfoot strike runners. J Sports Sci. 2021;39(1):78–83.

3. Fletcher JR, MacIntosh BR. Achilles tendon strain energy in distance running: consider the muscle energy cost. J Appl Physiol. 2015;118(2):193–9.

4. Ker RF, Bennett MB, Bibby SR, Kester RC, Alexander RM. The spring in the arch of the human foot. Nature. 1987;325(6100):147–9.

5. Lichtwark GA, Bougoulias K, Wilson AM. Muscle fascicle and series elastic element length changes along the length of the human gastrocnemius during walking and running. J Biomech. 2007;40(1):157–64.

6. Monte A, Maganaris C, Baltzopoulos V, Zamparo P. The influence of Achilles tendon mechanical behaviour on “apparent” efficiency during running at different speeds. Eur J Appl Physiol. 2020;120(11):2495–505.

7. Alexander RM. Energy-saving mechanisms in walking and running. J Exp Biol. 1991;160:55–69.

8. Bramble DM, Lieberman DE. Endurance running and the evolution of Homo. Nature. 2004;432(7015):345–52.

9. Cavagna GA, Kaneko M. Mechanical work and efficiency in level walking and running. J Physiol. 1977;268(2):467–81.

10. Charles JP, Suntaxi F, Anderst WJ. In vivo human lower limb muscle architecture dataset obtained using diffusion tensor imaging. PLOS ONE. 2019;14(10):e0223531.

11. Fletcher JR, Macintosh BR. Running Economy from a Muscle Energetics Perspective. Front Physiol. 2017;8:433.

12. Sanno M, Willwacher S, Epro G, Brüggemann G-P. Positive Work Contribution Shifts from Distal to Proximal Joints during a Prolonged Run. Med Sci Sports Exerc. 2018;50(12):2507–17.

13. Willer J, Allen SJ, Burden RJ, Folland JP. Neuromechanics of Middle-Distance Running Fatigue: A Key Role of the Plantarflexors? Med Sci Sports Exerc. 2021;53(10):2119–30.

14. Cigoja S, Fletcher JR, Nigg BM. Can changes in midsole bending stiffness of shoes affect the onset of joint work redistribution during a prolonged run? J Sport Health Sci. 2022;11(3):293–302.

15. Xu F, Montgomery DL. Effect of Prolonged Exercise at 65 and 80 % of VO2max on Running Economy. Int J Sports Med. 1995;16(5):309–13.

16. Sproule J. Running economy deteriorates following 60 min of exercise at 80% VO2max. Eur J Appl Physiol Occup Physiol. 1998;77:366–71.

17. Unhjem RJ. Changes in running economy and attainable maximal oxygen consumption in response to prolonged running: The impact of training status. Scand J Med Sci Sports. 2024;34(5):e14637.

18. Zavorsky GS, Montgomery DL, Pearsall DJ. Effect of intense interval workouts on running economy using three recovery durations. Eur J Appl Physiol Occup Physiol. 1998;77(3):224–30.

19. Zanini M, Folland JP, Blagrove RC. The Effect of 90 and 120 Min of Running on the Determinants of Endurance Performance in Well-Trained Male Marathon Runners. Scand J Med Sci Sports. 2025;35(5):e70076.

20. Sanno M, Epro G, Brüggemann G-P, Willwacher S. Running into Fatigue: The Effects of Footwear on Kinematics, Kinetics, and Energetics. Med Sci Sports Exerc. 2021;53(6):1217–27.

21. Enoka RM, Stuart DG. Neurobiology of muscle fatigue. J Appl Physiol. 1992;72(5):1631–48.

22. Hoogkamer W, Kipp S, Kram R. The Biomechanics of Competitive Male Runners in Three Marathon Racing Shoes: A Randomized Crossover Study. Sports Med. 2019;49(1):133–43.

23. Matijevich ES, Honert EC, Yang F, Lam W-K, Nigg BM. Greater foot and footwear mechanical work associated with less ankle joint work during running. Sports Biomech. 2025;24(6):1495–513.

24. Bermon S, Garrandes F, Szabo A, Berkovics I, Adami PE. Effect of Advanced Shoe Technology on the Evolution of Road Race Times in Male and Female Elite Runners. Front Sports Act Living. 2021;3:653173.

25. Willwacher S, Mai P, Helwig J, Hipper M, Utku B, Robbin J. Does Advanced Footwear Technology Improve Track and Road Racing Performance? An Explorative Analysis Based on the 100 Best Yearly Performances in the World Between 2010 and 2022. Sports Med Open. 2024;10(1):14.

26. Finni T, Kyröläinen H, Avela J, Komi P. Maximal but not submaximal performance is reduced by constant-speed 10-km run. J Sports Med Phys Fitness. 2004;43(4):411–7.

27. Girard O, Millet G, Jean-Paul M, Racinais S. Alteration in neuromuscular function after a 5 km running time trial. Eur J Appl Physiol. 2011;112:2323–30.

28. Nicol C, Kuitunen S, Kyröläinen H, Avela J, Komi PV. Effects of long- and short-term fatiguing stretch-shortening cycle exercises on reflex EMG and force of the tendon-muscle complex. Eur J Appl Physiol. 2003;90(5):470–9.

29. Hébert-Losier K, Finlayson SJ, Driller MW, Dubois B, Esculier J-F, Beaven CM. Metabolic and performance responses of male runners wearing 3 types of footwear: Nike Vaporfly 4%, Saucony Endorphin racing flats, and their own shoes. J Sport Health Sci. 2022;11(3):275–84.

30. Hoogkamer W, Kipp S, Frank JH, Farina EM, Luo G, Kram R. A Comparison of the Energetic Cost of Running in Marathon Racing Shoes. Sports Med. 2018;48(4):1009–19.

31. Hunter I, McLeod A, Valentine D, Low T, Ward J, Hager R. Running economy, mechanics, and marathon racing shoes. J Sports Sci. 2019;37(20):2367–73.

32. Farris DJ, Sawicki GS. The mechanics and energetics of human walking and running: a joint level perspective. J R Soc Interface. 2011;9(66):110–8.

33. Abbott BC, Bigland B, Ritchie JM. The physiological cost of negative work. J Physiol. 1952;117(3):380–90.

34. Margaria R. Positive and negative work performances and their efficiencies in human locomotion. Int Z Angew Physiol Einschl Arbeitsphysiol. 1968;25(4):339–51.

35. Glace BW, McHugh MP, Gleim GW. Effects of a 2-Hour Run on Metabolic Economy and Lower Extremity Strength in Men and Women. J Orthop Sports Phys Ther. 1998;27(3):189–96.

36. Hicks AL, Kent-Braun J, Ditor DS. Sex Differences in Human Skeletal Muscle Fatigue. Exerc Sport Sci Rev. 2001;29(3):109–12.

37. Temesi J, Arnal PJ, Rupp T, et al. Are Females More Resistant to Extreme Neuromuscular Fatigue? Med Sci Sports Exerc. 2015;47(7):1372–82.

38. Pallavi L, D Souza UJ, Shivaprakash G. Assessment of Musculoskeletal Strength and Levels of Fatigue during Different Phases of Menstrual Cycle in Young Adults. J Clin Diagn Res. 2017;11(2):CC11–3.

39. Meignié A, Duclos M, Carling C, et al. The Effects of Menstrual Cycle Phase on Elite Athlete Performance: A Critical and Systematic Review. Front Physiol. 2021;12:654585.

40. van Melick N, Meddeler BM, Hoogeboom TJ, Nijhuis-van der Sanden MWG, van Cingel REH. How to determine leg dominance: The agreement between self-reported and observed performance in healthy adults. PLoS One. 2017;12(12):e0189876.

41. Running Warehouse. Brooks Hyperion Tempo Shoe review. 2020; [cited 2025 Apr 18] Available from: https://www.runningwarehouse.com.au/Reviews/Brooks-Shoe-Reviews/brooks-hyperion-tempo.html?srsltid=AfmBOoo2a_rqbDwpWTomLAUnAFYkEKdcAm1GDSJ_MTzfqoqC2frI2Yv1.

42. Running Warehouse. ASICS METASPEED Sky+ Shoe Review. 2022; [cited 2025 Apr 18] Available from: https://www.runningwarehouse.com.au/reviews/ASICS-Shoe-reviews/metaspeed-sky-plus.html?srsltid=AfmBOop0QYPNKRfktNh59xsVtD_499HdwmWCOQlvdSjSWo7ZwPGFxNUb.

43. ASICS. METASPEED SKY+. 2022. [cited 2025 Feb 23] Available from: https://www.asics.com/au/en-au/metaspeed-sky%2B/p/AOP_1013A115-001.html.

44. Brooks. Hyperion Tempo. 2020; [cited 2025 Feb 23] Available from: https://www.brooksrunning.com.au/hyperion-tempo-mens-neutral-running-shoe/110339.html?colour=064&width=1D.

45. Willwacher S, Kurz M, Menne C, Schrödter E, Brüggemann G-P. Biomechanical response to altered footwear longitudinal bending stiffness in the early acceleration phase of sprinting. Footwear sci. 2016;8(2):99–108.

46. Binboğa E, Tok S, Catikkas F, Güven Ş, Dane S. The effects of verbal encouragement and conscientiousness on maximal voluntary contraction of the triceps surae muscle in elite athletes. J Sports Sci. 2013;31(9):982–988.

47. Place N, Lepers R, Deley G, Millet GY. Time Course of Neuromuscular Alterations during a Prolonged Running Exercise. Med Sci Sports Exerc. 2004;36(8):1347.

48. Saldanha A, Nordlund Ekblom MM, Thorstensson A. Central fatigue affects plantar flexor strength after prolonged running. Scand J Med Sci Sports. 2008;18(3):383–8.

49. Froyd C, Millet GY, Noakes TD. The development of peripheral fatigue and short-term recovery during self-paced high-intensity exercise. J Physiol. 2013;591(5):1339–46.

50. Hébert-Losier K, Fernandez MaR, Athens J, Kubo M, O’Neill S. A randomised crossover trial on the effects of foot starting position on calf raise test outcomes: Position does matter. Foot. 2024;60:102112.

51. Hébert-Losier K, Fernandez MR, Athens J, Kubo M, O’Neill S. A Randomized Crossover Trial on the Effects of Cadence on Calf Raise Test Outcomes: Cadence Does Matter. J Appl Biomech. 2025;41(2):179–88.

52. de Leva P. Adjustments to Zatsiorsky-Seluyanov’s segment inertia parameters. J Biomech. 1996;29(9):1223–30.

53. Bell AL, Brand RA, Pedersen DR. Prediction of hip joint centre location from external landmarks. Hum Mov Sci. 1989;8(1):3–16.

54. Hof AL. An explicit expression for the moment in multibody systems. J Biomech. 1992;25(10):1209–11.

55. Zelik KE, Honert EC. Ankle and foot power in gait analysis: Implications for science, technology and clinical assessment. J Biomech. 2018;75:1–12.

56. Bates D, Mächler M, Bolker B, Walker S. Fitting Linear Mixed-Effects Models Using lme4. J Stat Softw. 2015;67:1–48.

57. Lenth RV. emmeans: Estimated Marginal Means, aka Least-Squares Means. 2017;1.11.1. [cited 2025 Jun 16] Available from: https://CRAN.R-project.org/package=emmeans.

58. Delabastita T, Hollville E, Catteau A, Cortvriendt P, De Groote F, Vanwanseele B. Distal-to-proximal joint mechanics redistribution is a main contributor to reduced walking economy in older adults. Scand J Med Sci Sports. 2021;31(5):1036–47.

59. Hill AV. The Physiological Basis of Athletic Records1. Nature. 1925;116(2919):544–8.

60. Jones AM. The fourth dimension: physiological resilience as an independent determinant of endurance exercise performance. J Physiol. 2024;602(17):4113–28.

61. Zanini M, Folland JP, Wu H, Blagrove RC. Strength Training Improves Running Economy Durability and Fatigued High-Intensity Performance in Well-Trained Male Runners: A Randomized Control Trial. Med Sci Sports Exerc. 2025;57(7):1546.

62. Perrin TP, Gerey R, Morio CYM, et al. Effect of Footwear Longitudinal Bending Stiffness on Energy Cost, Biomechanics, and Fatigue during a Treadmill Half-Marathon. Med Sci Sports Exerc. 2025;57(4):657–67.

63. Nicol C, Avela J, Komi PV. The Stretch-Shortening Cycle. Sports Med. 2006;36(11):977–99.

64. Todd G, Gorman RB, Gandevia SC. Measurement and reproducibility of strength and voluntary activation of lower-limb muscles. Muscle Nerve. 2004;29(6):834–42.

65. Krüger RL, Aboodarda SJ, Jaimes LM, MacIntosh BR, Samozino P, Millet GY. Fatigue and recovery measured with dynamic properties versus isometric force: effects of exercise intensity. J Exp Biol. 2019;222(Pt 9):jeb197483.

66. James LP, Weakley J, Comfort P, Huynh M. The Relationship Between Isometric and Dynamic Strength Following Resistance Training: A Systematic Review, Meta-Analysis, and Level of Agreement. Int J Sports Physiol Perform. 2024;19(1):2–12.

67. Bohm S, Mersmann F, Schroll A, Arampatzis A. Speed-specific optimal contractile conditions of the human soleus muscle from slow to maximum running speed. J Exp Biol. 2023;226(22):jeb246437.

68. Monte A, Baltzopoulos V, Maganaris CN, Zamparo P. Gastrocnemius Medialis and Vastus Lateralis in vivo muscle-tendon behavior during running at increasing speeds. Scand J Med Sci Sports. 2020;30(7):1163–76.

