## Supplemental Table 1 for "Positive joint work redistribution in running: the role of plantar flexor fatigue and the effect of advanced footwear technology"

**Table 1** Individual participant details. $n$ = 12

| Participant | Age (yr) | Mass (kg) | Height (m) | Limb length (m) | Dominant limb | 10 km race time (mm:ss) | Testing run velocity (m $s^{-1}$) | Shoe size (US) | First shoe tested |
|---|---|---|---|---|---|---|---|---|---|
| 1 | 25 | 73.8 | 1.87 | 0.950 | Right | 41:00 | 4.00 | 11 | *AFT* |
| 2 | 25 | 87.8 | 1.89 | 0.970 | Right | 41:25 | 4.00 | 12 | *Traditional* |
| 3 | 24 | 78.0 | 1.88 | 0.957 | Right | 42:00 | 3.97 | 12 | *Traditional* |
| 4 | 21 | 78.0 | 1.84 | 0.954 | Right | 37:00 | 4.50 | 10 | *AFT* |
| 5 | 21 | 76.8 | 1.86 | 0.947 | Right | 40:00 | 4.17 | 10 | *Traditional* |
| 6 | 30 | 77.4 | 1.77 | 0.898 | Right | 37:00 | 4.50 | 10 | *AFT* |
| 7 | 34 | 74.2 | 1.84 | 0.916 | Right | 37:00 | 4.50 | 10 | *Traditional* |
| 8 | 31 | 72.4 | 1.78 | 0.881 | Right | 37:52 | 4.40 | 12 | *AFT* |
| 9 | 22 | 71.1 | 1.81 | 0.925 | Right | 37:00 | 4.50 | 11 | *Traditional* |
| 10 | 21 | 79.5 | 1.83 | 0.950 | Right | 38:20 | 4.35 | 12 | *AFT* |
| 11 | 26 | 83.0 | 1.88 | 0.900 | Right | 45:00 | 3.70 | 9 | *Traditional* |
| 12 | 22 | 74.3 | 1.79 | 0.920 | Left | 40:00 | 4.17 | 11 | *AFT* |
